# Cold Atmospheric Plasma induces silver nanoparticle uptake, oxidative dissolution and enhanced cytotoxicity in Glioblastoma multiforme cells

**DOI:** 10.1101/2020.02.28.969758

**Authors:** Eline Manaloto, Aoife Gowen, Anna Lesniak, Zhonglei He, Alan Casey, Patrick J Cullen, James F Curtin

**Affiliations:** BioPlasma Research Group, School of Food Science and Environmental Health, Technological University Dublin, Ireland; FOCAS Research Institute, Technological University Dublin, Ireland; Environmental Sustainability and Health Institute, Technological University Dublin, Ireland; UCD School of Biosystems and Food Engineering, UCD, Ireland; School of Physics and Clinical and Optometric Sciences, Technological University Dublin, Ireland; School of Chemical and Biomolecular Engineering, University of Sydney, Australia

**Keywords:** Cancer, Glioblastoma Multiforme, Silver Nanoparticles, Cold Atmospheric Plasma, Synergy

## Abstract

Silver nanoparticles (AgNP) emerged as a promising reagent for cancer therapy with oxidative stress implicated in the toxicity. Meanwhile, studies reported cold atmospheric plasma (CAP) generation of reactive oxygen and nitrogen species has selectivity towards cancer cells. Gold nanoparticles display synergistic cytotoxicity when combined with CAP against cancer cells but there is a paucity of information using AgNP, prompting to investigate the combined effects of CAP using dielectric barrier discharge system (voltage of 75 kV, current is 62.5mA, duty cycle of 7.5kVA and input frequency of 50-60Hz) and 10nm PVA-coated AgNP using U373MG Glioblastoma Multiforme cells. Cytotoxicity in U373MG cells was >100-fold greater when treated with both CAP and PVA-AgNP compared with either therapy alone (IC_50_ of 4.30 μg/mL with PVA-AgNP alone compared with 0.07 μg/mL after 25s CAP and 0.01 μg/mL 40s CAP). Combined cytotoxicity was ROS-dependent and was prevented using N-Acetyl Cysteine. A novel darkfield spectral imaging method investigated and quantified AgNP uptake in cells determining significantly enhanced uptake, aggregation and subcellular accumulation following CAP treatment, which was confirmed and quantified using atomic absorption spectroscopy. The results indicate that CAP decreases nanoparticle size, decreases surface charge distribution of AgNP and induces uptake, aggregation and enhanced cytotoxicity *in vitro*.

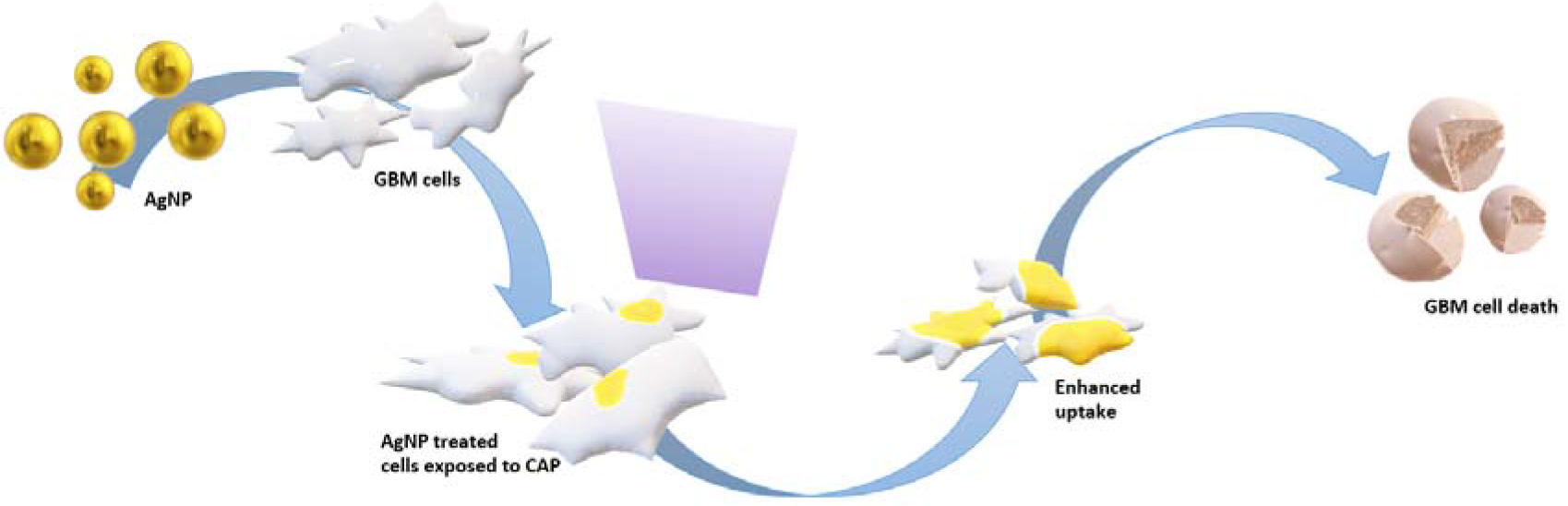

## Introduction

Glioblastoma Multiforme (GBM) is an aggressive grade IV astrocytoma. It is the most dominant form of central nervous system malignancy, accounting for 47.1% of all tumours diagnosed in the CNS[1]. The current predominant treatment is surgical resection followed by radiotherapy and chemotherapy with Temozolomide[2]. However, conventional treatments are rarely successful and are plagued with poor target delivery, poor efficacy and systemic toxicity. Despite intensive therapeutic strategies and medical care, approximately 5% of GBM patients survive five years after diagnosis[1]. In addition, more than 90% of GBM patients reveal recurrence at or near the primary site after treatment, underlining the need for a new effective therapeutic approach[3].

Nanoparticles (NP) have been used for therapy in diverse fields as a radiosensitiser[4], fluorescent labels[5], transfection vectors[6] and as drug carriers[7]. The size, shape and material composition confer advantageous properties on NP for various applications[8]. In particular, silver nanoparticles (AgNP) are the most widely used nanomaterial in consumer products such as household, cosmetics and healthcare-related products due to its antimicrobial properties through the release of silver ions[9]. The generation of reactive oxygen species has been associated with AgNP toxicity, making it useful for anticancer therapy[10]. Studies have shown that AgNP induce alterations in metabolic activity, cell morphology and decreased cell viability. Treatment with AgNP showed higher selectivity towards aggressive brain cancer human glioblastoma cells (U251) compared with a normal human lung fibroblast cells (IMR-90), leading to mitochondrial damage and increased in reactive oxygen production, resulting with DNA damage[11]. The combined NPs advantages of small size and large surface area has led to their use as drug delivery systems. In particular, AgNPs have attracted attention due to their intrinsic anticancer activity and effective drug delivery agents in previous studies[12–14]. Studies have shown AgNPs capability of crossing the blood brain barrier (BBB), providing opportunities and tackling challenges associated with NP drug delivery to the central nervous system (CNS)[15–17]. Recently, metal nanoparticles have been utilised to enhance cytotoxicity in cancer cells using oxidising treatments such as radiation therapy[18,19]. Liu, *et al,* demonstrated AgNP outperforming gold nanoparticles (AuNP) in radiosensitising glioma cell line U251, where combination of AgNP and radiotherapy showed significant anti-glioma effect *in vitro* and *in vivo* in comparison to AuNP[4]. These studies resulted to multitude applications of employing NPs to cancer treatments through a vast number of strategies.

Plasma treatment has shown potential as a future cancer therapy. Plasma is the fourth state of matter next to solid, liquid and gas. It can be artificially produced for its versatile applications[20]. Plasmas are classified as either thermal or non-thermal, also known as cold atmospheric plasma (CAP). The non-thermal nature of CAP coupled with a wide range of biological effects has led to the emergence of CAP across a range of biomedical applications including wound healing, dentistry and sterilisation[21]. CAP has low power requirements and is achieved at low or atmospheric pressure. CAP provides a rich environment of reactive oxygen species (ROS) such as singlet oxygen (O_2_), superoxide (O_2_^-^), ozone (O_3_), hydroxyl radicals (OH^-^), hydrogen peroxide (H_2_O_2_) and generates reactive nitrogen species (RNS) such as nitric oxide (NO) nitrite and nitrate anions (NO_2_ and NO_3_)[22]. Recent studies have shown CAP’s potential application in cancer therapy with biochemical features of cancer cells including high levels of ROS due to oncogenic transformation and with the application of CAP triggered self-perpetuating process of RONS induction, which effectively showed to induce apoptosis selectively against cancer cells overcoming the problem with conventional treatments[23]. In contrast to NPs systemic application, the effects of CAP are mostly associated with the location of treatment with few systemic effects observed and hence the recent trend in research of CAP is the interaction at cellular level [24]. The localised interaction of CAP with mouse fibroblast cells, BEL-7402 liver cancer cells and PAM212 cancer cells demonstrated detachment from extracellular matrix when treated[25]. CAP’s ability to change biochemical signalling intracellularly without thermal and electrical damage creates a suitable biomedical application[26]. The operating system of plasma discharged used in this study is the dielectric barrier discharge (DBD), DIT 120, which generates high voltage output of non-thermal plasma between two aluminium electrodes[27]. CAP induced by the DIT 120 system has previously been reported to induce cell death at higher exposures and enhance uptake of gold nanoparticles at lower exposures in U373MG glioblastoma multiforme cancer cells, the cell line used in this study[28,29].

A synergistic cytotoxic effect has been reported when CAP and various nanoparticles are combined, as first reported in 2009 by Kim, *et al.,* who found a 5-fold increase in cytotoxicity on G361 melanoma cancer skin cells when treated with ambient air CAP combined with antibody-conjugated AuNP [30]. Since then, studies using nanomaterials with various sizes and compositions have been used. For example, electrosprayed core-shell nanoparticle fabricated with 5-Fluorouracil synergistically inhibited cell growth of epithelial breast cancer cells MBA-MD-231 when used with CAP[31]. The plasma jet device using helium and oxygen gas in combination with Iron NPs significantly decreased viability of human breast adenocarcinoma cancer cells, MCF-7[32]. Our own group unveiled 25-fold enhanced cytotoxicity on U373MG cells when 20 nm citrate-capped AuNP were combined with non-toxic doses of CAP, demonstrating enhanced AuNP endocytosis and subcellular trafficking[29]. Meanwhile, interest in AgNP has shifted beyond antimicrobial use to potential additional anticancer applications[33–35]. Evidence is emerging that oxidative stress induced by low dose AgNP is implicated in their cytotoxicity[36–39]. Despite AgNP being the main commercial nanomaterial used worldwide, there are limited reports on combining AgNP with other current therapies to investigate its possible enhanced effect in comparison to various type of nanoparticles studied.

In consideration of the advantages of both AgNP and CAP, the importance of oxidative stress in both modes of cytotoxicity, the enhancement of AgNP toxicity when combined with other oxidising treatments and the finding that CAP can induce cellular uptake of nanomaterials, we chose to investigate whether synergistic cytotoxicity exists between AgNP combined with CAP and to explore the interaction using the U373MG GBM cell line model.

## Methods

### Chemicals

All chemicals used were obtained from Sigma-Aldrich (Vale Road, Arklow, County Wicklow, Ireland) unless specified otherwise.

### Cell culture

Human Glioblastoma Multiforme cells (U373MG-CD14) derived from a malignant tumour were obtained from Dr. Michael McCarthy (Trinity College Dublin). U373MG cells were maintained in DMEM/F-12 Ham (Sigma-Aldrich, D8062) and were supplemented with 10% v/v foetal bovine serum. The cell line was incubated at 37°C and 100% humidity containing 5% CO_2_. U373MG were sub-cultured at 70-80% confluency using 1:1 ratio of 0.25% trypsin (Gibco by Life Technologies, UK) in Hank’s balanced salt solution and 0.1% EDTA in phosphate buffer saline.

### AgNP preparation

The top-down synthesis used in the study was previously reported by Mavani *et al*[40]. Cold synthesis was employed with chemical reduction of silver nitrate (AgNO_3_) of 0.001M with icecold reducing agent sodium borohydride (NABH_4_) of 0.002M in the presence of a stabiliser formed AgNP. 2ml of 1% polyvinyl alcohol (PVA) in millipore water (Simplicity 185, 18.2 MΩ.cm at 25°C resistivity) was added with 2ml of silver nitrate and mixed well[41]. The icecold reducing agent was stirred for 20 minutes and 2 ml of mixed AgNO_3_PVA were added 1 drop per second approximately and reaction was stopped. PVA stabilised silver nanoparticles (PVA-AgNP) mixture was concentrated by using an ultrafiltration tube of 3kDa (Sartorius, UK) with centrifuge, Heraeus Megafuge 16R (Thermo Scientific) at 5000 rpm. PVA-AgNP were stored away from direct exposure to light at 4°C. The chemical reduction of AgNO_3_ can be written as:

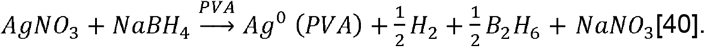

### AgNP characterisation

The primary characterisation of synthesised PVA-AgNP was achieved by Ultra-Violet Spectroscopy (UV-Vis) (Shimadzu, UV-1800) and the stability of PVA-AgNP prepared were observed for 6 months. Dynamic Light Scattering (DLS) was used to measure the hydrodynamic diameter (D_h_) and zeta potential of PVA-AgNP (Malvern Instruments, UK). The size, shape and morphology of PVA-AgNP was assessed by scanning transmission electron microscopy (STEM) (Hitachi SU 6600) with 3μl of PVA-AgNP aqueous solution placed on a carbon-coated copper grid and the samples were allowed to dry to obtain the highest yield of particles on the grid. PVA-AgNP concentration measurements were performed by atomic absorption spectroscopy (AAS) using (Varian SpectrAA 200) against Ag calibration standard at 1, 2, 3, 4, 5 μg Ag/ml prepared from 1 g/L standard. AgNP samples were prepared by 1/10 dilution by adding 1ml concentrated PVA-AgNP to 9 mL of millipore water.

### Cold atmospheric plasma device

The CAP system used was a dielectric barrier discharge (DBD) device. It is a novel prototype atmospheric low temperature plasma generator[27]. The CAP-DBD is composed of a variable high voltage transformer with an input voltage of 230 V at 50 Hz and a maximum high voltage output of 120kV. It consists of two 15-cm-diameter aluminium disc electrodes, separated by 26.6 mm and a 1.2mm thick polypropylene sheet which is used as a dielectric barrier and platform for holding cell samples. The voltage was monitored using an InfiniVision 2000 X-Series Oscilloscope (Agilent Technologies Inc., Santa Clara, CA, USA). For U373MG exposure to CAP, culture media was removed prior to CAP exposure leaving 10 μl culture media in 96 well plates and 500 μl of culture media in 60mm petri dishes to avoid sample drying. Fresh culture media was added immediately after exposure to CAP with a total of 100 μl in 96 well plates and 3 ml in 60mm petri dishes. The samples were treated at 75 kV at different exposure times from 0-80s.

### Cell viability assay

Alamar Blue was used as a parameter for measuring cytotoxicity[28,42]. Alamar blue assay was analysed according to manufacturer’s protocol (Invitrogen by Thermo Fisher Scientific, USA). Firstly, U373MG cells (1×10^4^ cells per well) were seeded into 96-well plates and allowed to adhere overnight. After seeding, 100μl of various concentration with 1:3 serial dilution of PVA-AgNP in DMEM were added between 0 μg/ml and 17.54 μg/ml. 1M H_2_O_2_ was used as a positive control for all cell viability assays. After preloading with PVA-AgNP for 24 hours, the cells were then exposed to CAP from 0-80s at 75kV as outlined above. The plates were incubated for a total of 48h post CAP treatment at 37°C. Cells were then washed with PBS and 10% Alamar blue was added into the wells for 3h at 37°C. The fluorescence was measured using excitation wavelength of 530nm and emission wavelength of 590nm on plate reader (SpectraMax M3, Molecular Devices (UK) Ltd). The protective effect of N-Acetyl Cysteine (NAC) was evaluated by pre-treating 4mM NAC on U373MG cells for 1h followed by treatment of PVA-AgNP at stated concentrations for 24h and exposed to CAP at 0s, 25s and 40s for another 24h.

### Flow cytometry for live and dead cell staining

Propidium iodide (PI) was used to demonstrate live and dead cell staining with flow cytometer BD Accuri C6 (BD, Oxford, UK). U373MG cells were seeded in 6-well plate at 2.5 x 10^5^ cells per well and was incubated at 37°C overnight to allow adherence. Cells were treated with PVA-AgNP at low-nontoxic dose of 0.07μg/mL for 24h and were exposed to CAP at 75kV for another 24h. Cells were harvested including pre-existing media and centrifuge to form a pellet. The pelleted cells were resuspended in 1ml PBS and was stained with PI for 1 minute with concentration of 10μg/mL. PI fluorescence was detected using FL2 vs FSC demonstrating binding of PI to nuclear degradation from dead cells.

### Measurement of CAP effect on AgNP size

The effect of CAP on PVA-AgNP size was measured as follows: concentrated PVA-AgNP obtained from ultrafiltration to remove excess unwanted reactants including NaBH_4_ was resuspended in millipore water, in synthesis solution containing unwanted reactants and resuspending concentrated PVA-AgNP in 4mM NaBH_4_. The samples were exposed to CAP at different exposure times from 0-80s. The hydrodynamic size of the NPs was determined using DLS, Zetasizer Nano ZS. The morphology was determined using STEM (Hitachi SU 6600), as described above.

### Darkfield spectral imaging of AgNP uptake: image acquisition and uptake analysis

Optical observation of AgNP uptake was performed using darkfield spectral imaging (SI). The darkfield microscopy system utilised in this study consists of a darkfield microscope (BX51, Olympus, Ltd) coupled to a Vis-NIR spectrograph (ImSpector V10E, Specim Ltd) and a Pelier cooled CCD detector, combined with a metal halide light source, liquid light guide and Cytoviva optical illuminator (CytoViva Inc., Auburn, AL, USA) and a motorised XY stage controller Images were obtained using a 100X oil immersion objective and oil immersion swing condenser.

Detection and confirmation of AgNP uptake were determined by spectral mapping, using samples of fixed U373MG cells alone (38 images from slides), cells treated with AgNP (42 images from slides), cells treated with CAP (42 images from * slides) and cells treated with combined AgNP-CAP (50 images from slides). U373MG cells were fixed by 4% paraformaldehyde on cover glass slip mounted with Mowiol 4-88 onto a rectangular cover glass slide (Corning cover glass, Sigma-Aldrich). Images were obtained over a time period of 10 weeks as each image took approx. 15 minutes to obtain.

The spectral images were acquired using ENVI 4.8 software and analysed using MATLAB (MATLAB R2019a, The MathWorks Ltd) functions written in house and from the image processing toolbox.

False colour red-green-blue (RGB) images were constructed from the spectral images by assigning the following wavebands to each channel:

Red: calculate mean image of waveband channels from 516 – 552 nm and scale to intensity values between 0-2000.

Green: calculate mean image of waveband channels from 544 – 618 nm and scale to intensity values between 0-2000.

Blue: calculate mean image of waveband channels from 704 – 735 nm and scale to intensity values between 0-2000.

PCA was applied to the false colour images to create a mask between the cell and background (thresholding the first principal component score image such that all pixels with a values below 2 were set to one provided sufficient separation between cells and background). In some cases, multiple cells were attached to each other in the mask images. In order to be able to analyse each cell individually, these cells were separated manually by drawing a line across the narrowest connecting point in the mask image. In addition, some cells were only partially represented in a given field of view – such cells were removed from the analysis. Further, some pixels that were not identified as cells were highlighted in the mask – in order to remove them from the analysis, any regions with a pixel size smaller than 300 pixels were automatically removed from the analysis.

Standard normal variate pre-processing (i.e. subtracting the mean from each spectrum and dividing by the standard deviation) was applied to spectra and a partial least squares discriminant analysis model was developed to discriminate between NPs and cells. This model was subsequently applied to all images, facilitating identification of NPs from their spectra. Using the ‘regionprops’ function of the MATLAB image processing toolbox, the size and circularity of each cell and number and size of NPs per cell was calculated. The median number of NPs identified per cell was then calculated for each treatment.

### AAS measurement of AgNP uptake

Atomic Absorption Spectroscopy (AAS) was further employed to measure uptake of AgNP. U37MG cells were grown in petri dishes with cell density of 2.5 x10^6^ cells per dish. The samples were negative control with U373MG cells alone, cells treated with AgNP, cells exposed to CAP, the combined therapy on U373MG cells and positive control treated with 1mM H_2_O_2_. The cells were treated with AgNP for 24h and after with 25s CAP for another 24h. The cell samples were trypsinised and resuspended in the existing culture media. The samples were then analysed by AAS for AgNP detection. Triplicate readings were analysed for each sample and the results were expressed as the mean amount of Ag in pg/cell.

### Statistical analysis

The data of the experiments are expressed as mean ± standard deviation of replicates from at least three independent experiments, unless specified otherwise. Statistical analysis and curve fitting presented in results were completed using Prism 7 (GraphPad Software). Analysis of data distribution was performed using two-way ANOVA and three-way ANOVA where indicated to analyse the differences of significance between the control group and treated groups. The following P values were deemed statistically significant, **P<0.01, ***P<0.001, ****P<0.0001. Therapeutic synergism between PVA-AgNP and CAP was evaluated using isobologram analysis. CompuSyn software determined the combinational index (CI) where, CI>1 is antagonism, CI=1 is additive and CI<1 is synergism. Descriptive statistics (median, standard deviation) were calculated on cell size, circularity, NP size and number as derived from spectral imaging measurements.

### Data Availability

All datasets can be viewed in tabular form in Supplementary Information.

## Results

### Silver Nanoparticles Characterisation

PVA-AgNP were prepared and characterised as indicated in the methods section. STEM analysis of PVA-AgNP confirmed production of nanoparticles that are spherical in shape, approximately 10nm in size and well dispersed (see Figure 1a). The presence of a plasmon absorption band (400 nm), which is a main characteristic of AgNP, was evident and remained relatively unchanged over 6 months indicating the production of highly stable AgNP (see Figure 1b)[43]. The particle size range by DLS analysis was determined to be 8-12 nm with an average of 10.68 ± 1.98 nm diameter size on first synthesis and after 6 months of 9.59 ± 2.52 nm (see Figure 1c and 1d). PVA-AgNP polydispersity index (PDI) at first synthesis was 0.415 and after 6 months of 0.155, which is deemed acceptable with ISO standard document 22412:2008 with data remaining between the acceptable PDI range values of 0.05 to 0.7.

**Figure 1.**
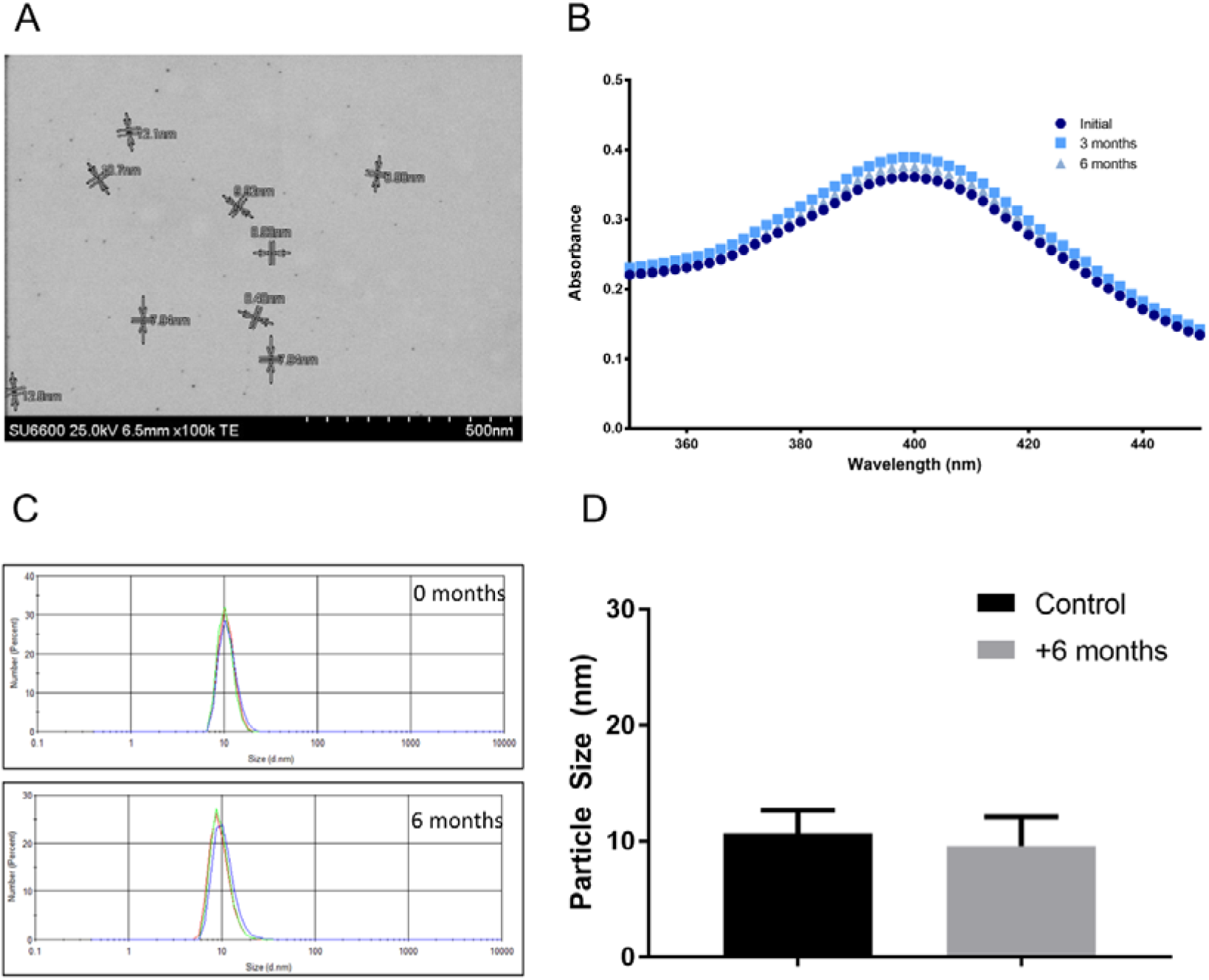
Characterisation of Silver Nanoparticles (AgNP). **(A)** Representative scanning transmission electron microscopy images of AgNP was spherical in shape and well dispersed in aqueous solution. **(B)** Representation of AgNP maximum absorbance by UV-Visible spectrum of stable aqueous AgNP mixture observed for 6 months. (**C, D**) Size distribution measurement of stable PVA-AgNP for six months by dynamic light scattering 10.68 ±1.98nm to 9.59 ±2.52nm.

### Cytotoxic Effect of AgNP in Combination with CAP on Cellular Viability

Cytotoxicity was examined using Alamar Blue and propidium iodide. U373MG cells were first treated with different concentrations of PVA-AgNPs (0-17.54 *μg/ml*) in combination with CAP (0-40s at 75kV) and incubated for 48 hours prior to measuring viability using Alamar blue (see Figure 2a). The cell viability decreased in a dose-dependent manner after AgNP treatment alone with IC_50_ of 4.730 μg/mL (95% confidence range from 3.094 to 6.084 μg/mL). From previous work with the same DBD prototype used in this study, it was determined that the IC_50_ with CAP treatment alone is 74.26s (95% confidence range of 47.24-116.8s) on U373MG cells[28]. In the current study, we combined treatment of AgNP with a range of low CAP exposures (i.e. 5s, 10s, 25s and 40s at 75kV). We found that cells displayed significant cytotoxicity with IC_50_ of 0.079 μg/mL and 0.01 μg/mL when treated with CAP for 25s and 40s respectively (95% confidence range from 0.0539 to 0.1139 μg/mL for 25s and 0.0069 to 0.0138 μg/mL for 40s) (see Figure 2a). Two-Way ANOVA with Tukey’s multiple comparison post hoc test in figure 2b confirmed significant decrease in IC_50_ values caused by 25s and 40s CAP exposure (see Figure 2c, **P<0.01, ***P<0.001, ****P<0.0001). The combined AgNP and CAP treatment exhibited 67-fold increase in cytotoxicity (IC_50_) when combined with 25s CAP and greater than 100-fold increase in cytotoxicity (IC_50_) with 40s CAP, which indicates a beneficial combinational anti-cancer therapy. This shows an enhanced toxic effect on U373MG cells when treated with combined therapy than with AgNP or CAP alone. No enhanced cytotoxicity was noted for 5s or 10 s doses with CAP in combination with AgNP, suggesting synergistic cytotoxicity only occurs between a minimum and maximum threshold dose for CAP. The isobologram on figure 2c determined the synergistic effect of PVA-AgNP combined with CAP resulting to combination index value (CI) of less than 1.00. The CI of PVA-AgNP with 25s CAP is 0.35 and 0.54 with 40s CAP. PVA-AgNP with shorter CAP exposure time resulted to CI of 0.93 and 0.96 with 5s CAP and 10s CAP respectively suggestive of an additive effect.

**Figure 2.**
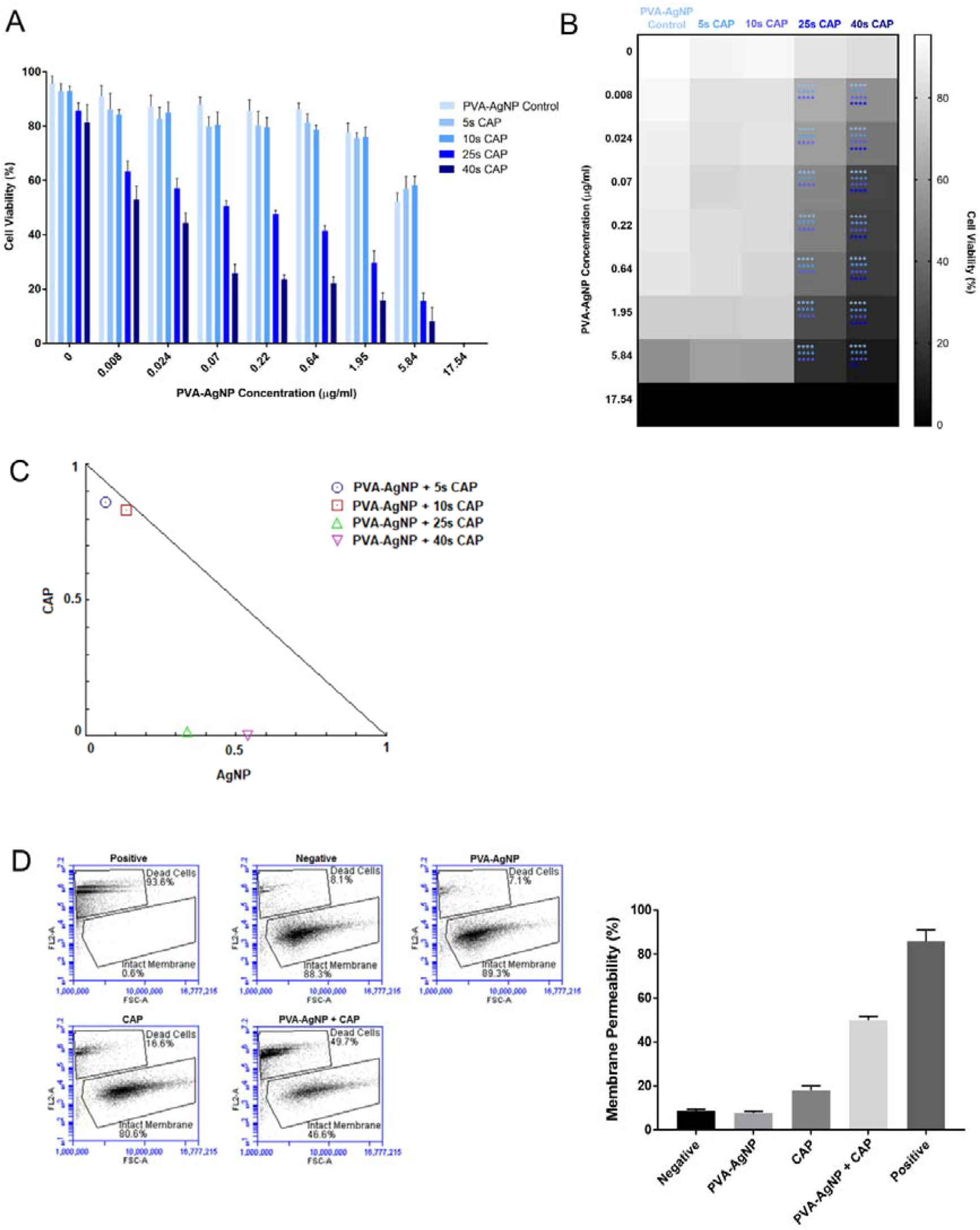
Synergistic Cytotoxicity of combined AgNP with Cold Atmospheric Plasma (CAP). **(A)** Combinational therapy effect of AgNP with CAP on U373MG cells. Cells were treated with different concentration of AgNP for 24h followed by different time exposure with CAP at 75kV. **(B)** Heat map representation of results from Figure 2A, showing the darkest region with the least viable cell to the lightest region with the most viable cell from combinational AgNP with CAP at different time exposure. Statistical analysis carried out using two-way ANOVA for Figure 2A, two-way ANOVA with Tukey’s multiple comparison post-test (**P<0.01, ***P<0.001, ****P<0.0001). All experiments were repeated at least three times. **(C)** Isobologram analysis of the combinational effect of AgNP-CAP. The single doses CAP on y-axis and AgNP on x-axis were used to draw the line of additivity. The localisation of combined AgNP-CAP at different time exposures can be translated to synergism CI<1, additivity CI=1 or antagonism C1>1. **(D)** Apoptotic nuclear membrane degradation was validated for the combined PVA-AgNP and CAP at 75kV using propidium iodide (10μg/ml) with flow cytometry. The represented data was normalised to the negative control and is displayed as %mean ± SD (n=3).

The enhanced cell death was confirmed and quantified by employing propidium iodide with flow cytometry. Figure 2d demonstrates the enhanced uptake of PI with the combined therapy of PVA-AgNP and CAP, in which PI has bound to the nucleic acids within the nucleus of AgNP-CAP treated GBM cells and compared to the negative untreated control. The size and granularity of dying cells changes, where dead cells are smaller and more granular resulting in high PI fluorescence in FL2 and live cells are larger and less granular resulting in low PI fluorescence as can be seen in the dot plot of figure 2d with FL2 in log scale vs. FSC in linear scale, which was used to distinguish live from dead GBM cells.

### AgNP-induced Cytotoxicity Protected by NAC

The main component of CAP is the generation of RONS and many have linked oxidative stress with AgNPs toxicity in previous studies[36–39]. NAC is a scavenger of oxygen-free radicals and directly interacts with reactive ionised species. The protective effect of NAC on AgNP and in combination with CAP can be seen in figure 3a. The IC_50_ value for AgNP control was 4.9 μg/ml with 95% confidence range of 3.759 to 6.081 μg/ml. The IC_50_ value for AgNP with pre-treated NAC was 6.57 μg/ml with 95% confidence range of 5.989 to 7.130 μg/ml. The IC_50_ value for the combinational AgNP-CAP without NAC at 25s was 0.06 μg/mL (95% confidence range of 0.0355 to 0.112 μg/mL), whereas a 10-fold decrease in toxicity was observed when cells were pre-treated with NAC with IC_50_ of 0.7 μg/mL (95% confidence range of 0.383 to 1.655 μg/mL). Three-Way ANOVA with Tukey’s multiple comparison post hoc test in figure 3b confirmed to be statistically significant (see figure 3b, **P<0.01, ***P<0.001, ****P<0.0001).

**Figure 3.**
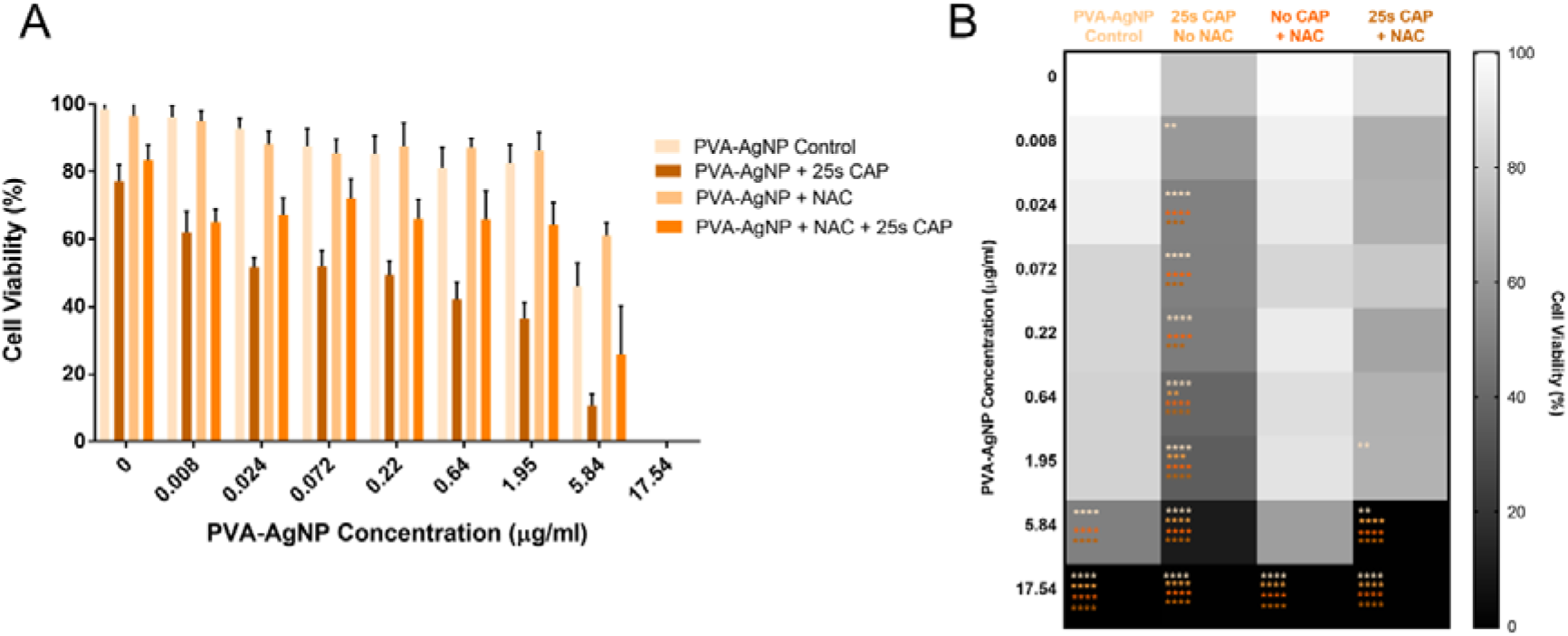
Protective effect of NAC. **(A)** Pre-treatment of N-Acetyl-L-cysteine (NAC) showed protective effects against AgNP combined with CAP after 48h using Alamar Blue assay. **(B)** Heat map interpretation of protective effects of NAC on combined therapy with the lighter the colour the more viable cells and the darker the heat map with less viable cells. Statistical analysis carried out on data presented in figure 3A using three-way ANOVA for 2C and 2D with Tukey’s multiple comparison posttest (**P<0.01, ***P<0.001, ****P<0.0001). All experiments were repeated at least three times.

### The effect of CAP on AgNP size

Our data indicate that ROS-dependent cytotoxicity is induced in GBM cells treated with a combination of CAP and AgNP. The toxicity is likely due to one or more of the following: cellular rate of uptake, increase in cell membrane permeability, changes to nanoparticle size and morphology, altered dispersion or agglomeration and rate of dissolution. It has been reported that NPs can be prepared using an electrical discharge[44–46]. We therefore investigated the effect of CAP on AgNP. 10nm freshly synthesised AgNP including unwanted reactants in the existing solution containing excess NaBH_4_ (4mM) were exposed to the DBD-CAP device (75kV, 0-80s). The size of AgNP significantly decreases in a dose-dependent manner when exposed to CAP (Figure 4a). In contrast to this, purified AgNP resuspended in fresh millipore water without unwanted reactants with CAP did not change significantly in size. We confirmed this using STEM. AgNP in a solution of the reductive agent, NaBH_4_ (4mM) showed decrease in size (5 nm) when exposed to 25s CAP compared with controls without CAP treatment (10 nm) (see Figure 4b). Nanoparticle morphology and dispersion did not change during CAP exposure, and AgNP remained spherical in shape, uniformly dispersed with no visible aggregation in the samples was observed.

**Figure 4.**
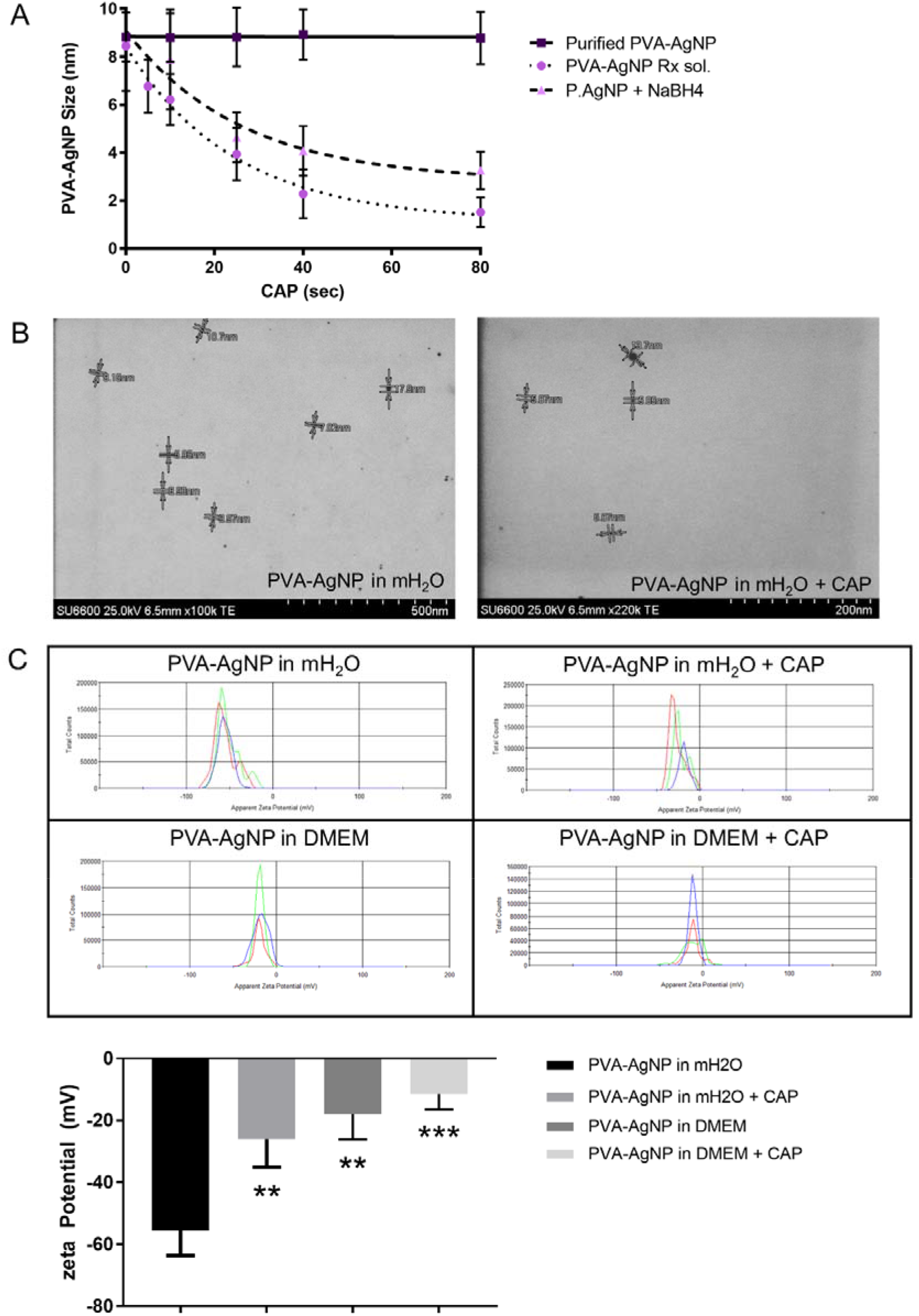
Comparative Results on the Effect of CAP on AgNP Physical Properties. **(A)** Overview of PVA-AgNP size change with different time exposure to CAP with and without presence of reductive agent. Data represented mean ± standard error of the mean (n=9). **(B)** STEM images displaying size variation of AgNP in aqueous solution without CAP exposure and 25s exposure with CAP at 75kV. **(C)** Zeta potential of AgNP before and after CAP treatment in millipore water (mH_2_O) and culture media (DMEM). Representation of zeta potential of AgNP before CAP treatment with −55.6 ±8.06mV to −26.8 ±9.08mV after 25s CAP exposure in mH_2_O and −18.0 ± 8.70mV without CAP exposure to 11.40 ± 5.06mV after CAP exposure at 75kV in culture media.

The recognised value for zeta potential that is anything higher than positive or negative 30mV is a stable suspension[47]. Figure 4c shows an electrostatically stabled nanoparticle suspension of −55.6 ± 8.08mV with conductivity of 0.360mS/cm and after CAP exposure of 25s the nanoparticle suspension has zeta potential of −26.80 ± 9.08mV with conductivity of 0.25mS/cm for PVA-AgNPs suspended in millipore water. In contrast to PVA-AgNP in millipore suspension, PVA-AgNPs in culture media (DMEM-F12) has changed the zeta potential value of −18.0 ± 8.70mV with conductivity of 0.011mS/cm and after CAP exposure of PVA-AgNP in DMEM-F12 the zeta potential decreased further to −11.40 ± 5.06mV with conductivity of 0.30mS/cm. The change of zeta potential of NPs was previously reported in other studies that affects the internalisation process during uptake of NPs by cells. Uptake of PVA-AgNP was further explored and can be seen in figure 5.

**Figure 5.**
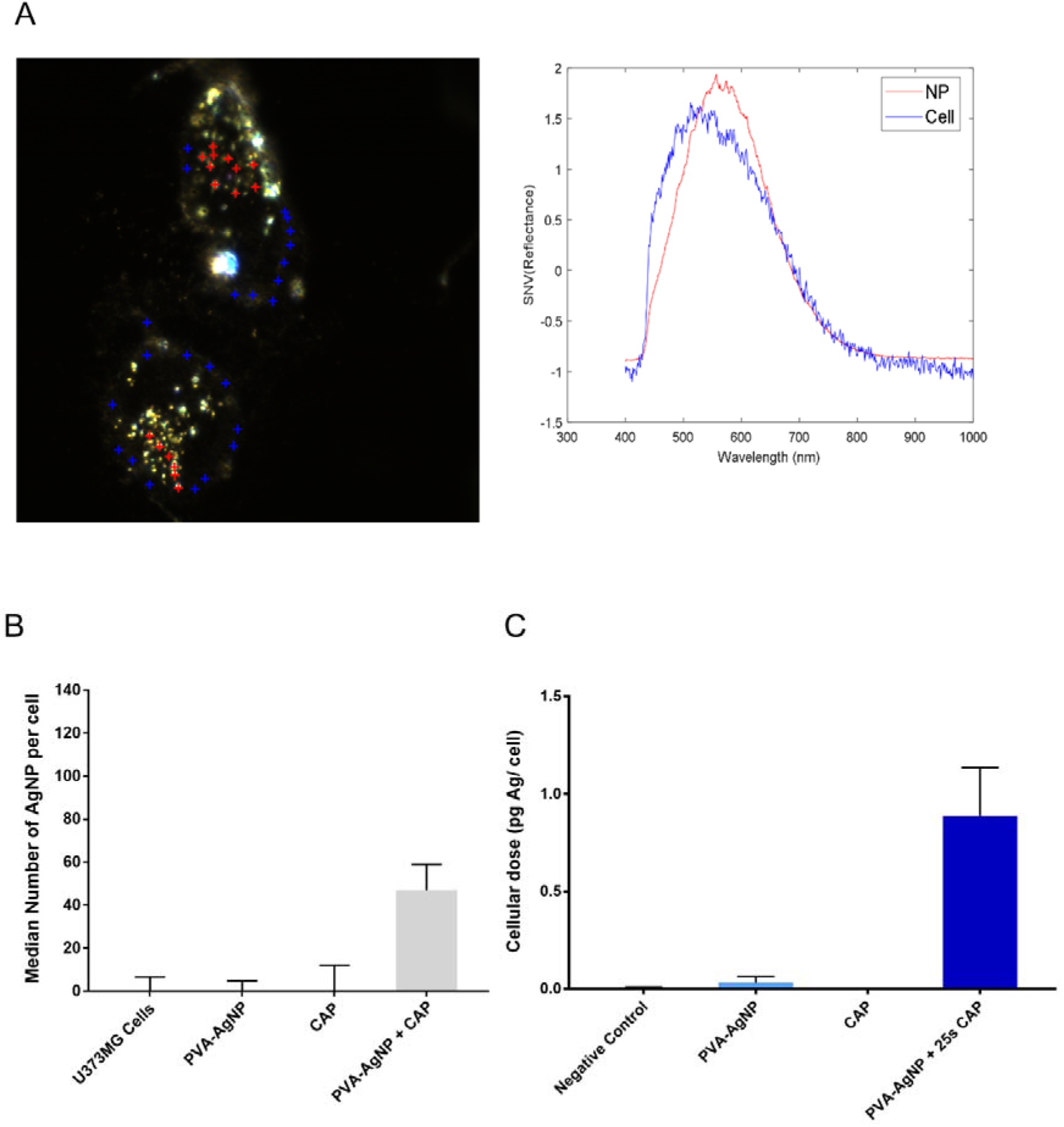
Uptake and Dissolution of AgNP by CAP. **(A)** Spectral imaging of mapped PVA-AgNP with red pixels in U373MG cells combined with CAP and representative spectral response of U373MG cells and PVA-AgNP in cells. **(B)** Representation of SI analysis on quantifying median number of AgNP/cell with only the combined therapy of AgNP-CAP resulted to 47 NPs per cell. Data shown was normalised to the negative untreated control and is shown as %median ± S.E.M. **(D)** AgNP uptake before and after CAP treatment was confirmed and quantified using AAS. The mean concentration of silver per cell was 0.030pg Ag/cell for PVA-AgNP treatment alone and 0.89pg Ag/cell for combined treatment of AgNP-CAP.

### Uptake of AgNP in U373MG cells

The study further investigated whether the significant difference on cytotoxicity of AgNP treated alone compared to combination of AgNP-CAP could be explained by differences in cellular uptake and localisation intracellularly. Visualisation of cell morphology, nanoparticle distribution and particle localisation after treatment with low dose of 0.07 μg/ml AgNP alone and in combination with 25s CAP at 75kV was investigated using Spectral imaging (SI) (see Figure 5a). SI offer quantitative analysis capturing both biological and materials at nanoscale level. The classification tools with dominant spectral signatures enables to characterise nanoparticles in populations and individually and morphology of cells prior and after treatment. Figure 5a shows images of *in-vitro* cells with false RGB setting derived from SI Cytoviva and a comparison mathematical modelling analysed from Matlab viewing cell morphology of brain cancer cells U373MG without NP treatment showing a healthy astrocyte-shaped cells, followed by U373MG remaining its healthy astrocyte-shape treated with low dose of PVA-AgNP present, cells treated with CAP for 25s showing cellular stress with morphological changes losing the astrocyte shape with no NPs evident and the combination therapy of AgNP-CAP presenting cell disruption and increase of PVA-AgNP visible in the cells. The comparison of the mathematical with false-RGB images allows to visualise and localise PVA-AgNP present in U373MG cells when treated. A library of spectra was derived from the scattered light of cells and PVA-AgNP identifying unique spectral signatures that is highly repeatable in approx. 50 images per sample (see Figure 5a). The spectral profile is graphed as wavelength (nm) versus intensity of scattered light (a.u.) presenting spectral signatures and interaction of NPs to cells with spectral response showing a broad scattering spectrum of U373MG cells alone are at 520nm with low intensity, PVA-AgNPs spectral response in cells is at 520nm with higher intensity and an enhanced intensity of PVA-AgNP showing a shift of the resonance peak to 570nm presents NPs aggregating into larger sizes.

SI allows to quickly identify, map PVA-AgNP present in cells and provide class distribution confirming the total number and size of NPs in cell. Figure 5b displays NPs class distribution with only the combination therapy of AgNP-CAP detecting median number of 47 NPs per cell with standard deviation of 84.42 NPs and a median NP size of 8nm with standard deviation of 9.7nm. However, numerical data derived from analysis of spectral images should be considered approximate, since light scattering from different cellular compartments and may hinder thorough viewing or may cause false positive detection of NPs. To further verify the uptake of AgNP as observed from SI, the cellular dose of AgNPs in U373MG cells was quantified using Atomic Absorption Spectroscopy. The mean concentration of Ag per cell was observed to be 0.030 pg Ag/ cell for AgNP treatment alone (95% confidence range of 0.022 to 0.083 pg Ag/ cell) and 0.89 pg Ag/ cell for combined therapy AgNP-CAP (95% confidence range of 0.50 to 1.28 pg Ag/ cell) (see Figure 2d). This correlates with our previous study (He *et al*) when we found that combination of gold nanoparticles with CAP increased nanoparticle uptake[29].

## Discussion

The advancement of nanotechnology and plasma medicine in biomedical applications tackle the same set of challenges but have been developed independently and often along different routes. The similarities and contrast of nanoparticles and cold atmospheric plasma have scoped their intriguing predetermined therapeutic use, particularly the selectivity against cancer cells with the selective application process of CAP as well as functionalising nanoparticles for targeted delivery to cancer cells, chemical reactivity through generation of ROS by CAP and AgNP, finally the safety to healthy cells and tissue. Concurrently, synergy captured from the combination of nanoparticle and cold atmospheric plasma with recent studies including gold nanoparticles, iron nanoparticles and drug-loaded core shell nanoparticles with cold atmospheric plasma have demonstrated highly promising benefits for medicine[23,27–29]. Despite the reports of CAP sensitivity to cancer cells, we and others have demonstrated that the Glioblastoma multiforme cell line U373MG is relatively resistant to CAP treatment[28] and approaches to overcome this inherent resistance will be necessary in a clinical setting. In addition, with the ever-growing interest of combined cancer therapy with nanotechnology and plasma medicine, the combination of AgNP with CAP has not yet been reported. In this study, the combined synergistic effect of PVA stabilised AgNP with CAP on U373MG was evaluated and the interaction between CAP and AgNPs physical properties and its effect on cell morphology were demonstrated.

The restrictive approach in this study focuses on strategic minimal dosing of AgNP in combination with CAP to achieve a targeted cytotoxic effect on Glioblastoma cell line U373MG. NPs physico-chemical characteristics and their effects play a major role on biological systems, where NPs are reported to penetrate and accumulate in different tissues showing high mobility. The shape, morphological structure and size are the principal parameters of NPs along with its chemical composition and its inherent toxicological properties[8,48]. Consequently, the characterisation of NPs physical properties is key to compare its biological and toxicological response. In this study, an in-house synthesised spherical AgNP stabilised with PVA with size 10.68 ± 1.98 nm showed high stability in aqueous solution for six months, which is in agreement with previous studies on size-controlled synthesis of AgNP stabilised with PVA[41,49]. Toxicological studies were implemented to determine *in vitro* synergistic cytotoxic effects of AgNP when combined with CAP. The results confirmed a dose-dependent reduction in cell viability with the control group (PVA-AgNP alone treated U373MG cells) with IC_50_ of 4.74 μg/ml. Many studies have shown AgNPs cytotoxic effect on variety of cancer cells, with all noticeably reporting AgNP inducing dose-dependent cytotoxicity including DNA damage and oxidative stress resulting in cell death[36–39]. Interestingly as viewed in figure 2a, the combination of AgNP with CAP increased more than 100-fold of cytotoxicity in comparison to AgNP treatment alone with IC_50_ of AgNP combined with 25s CAP at 75kV is 0.077 μg/ml and when combined with 40s CAP at 75kV, IC_50_ was 0.0087 μg/ml. CAP similarly generates ROS and can be localised by selective application process at a milli or micro scale to cancer cells[50,51]. Consistent with the findings regarding to oxidative stress correlation to cell death with AgNP and CAP continuously mounts the evidence to show that generation of ROS is highly related to the mechanism of AgNP and CAP with the results in current study showed that the cytotoxicity induced by AgNP alone, as well as the combinational treatment AgNP-CAP were efficiently prevented prior NAC treatment up to 10-fold (see Figure 3). The results determine that oxidative stress is responsible for the cytotoxicity of AgNPs and CAP, which is compliant with previous studies portraying the protective effect of NAC when treated with either AgNP or CAP resulting to recovery of proliferative cells[52–55].

Tseng *et al,* has previously reported metal nanoparticle fabrication using CAP in the form of electric discharge machine, where the generation of arc discharge between two electrodes disintegrate silver rod in liquid producing silver nanoparticles[45,56,57]. Due to this phenomenon, the current study next investigated the effects of CAP on AgNP physical properties. CAP evidently reduced the diameter size of AgNP with longer exposure time in presence of reducing agent (see Figure 4a). A study has investigated hydrolysis of NaBH_4_ in aqueous solution, it was reported that the stability of NaBH_4_ decreases with elevated temperature[58–60]. It can be hypothesised with the findings that CAP’s effect on AgNP leads to generation of discharge between two electrodes producing a thermal effect on AgNP solution with borohydride present, this increases temperature and thus accelerates hydrolysis reaction. Furthermore, CAP’s alteration on PVA-AgNP size in our study resulted in altered electronic properties on the surface of NP (see figure 4C). Several studies have shown size-dependent effects on cytotoxicity using silver nanoparticles[61–63]. In many cases, the levels of cell death are increased when smaller nanoparticles are used, and this is believed to be due to a larger surface area and enhanced rates of endocytosis. However, the effect of size on cytotoxicity in these studies is relatively small (approx. 2-5 fold) and unlikely to be solely responsible for the synergistic cytotoxicity between CAP and AgNP observed in our study. Many reports stated the standard stable nanoparticle suspension is anything higher than positive or negative 30mV. The higher value of positive or negative zeta potential has been studied to show in nature to repel each other and not come together. The particles tend to aggregate and flocculate with lower zeta potential values due to the absence of repulsive forces that hinders agglomeration[64–66]. Studies have reported that the greater the negative charge the less toxic the nanoparticles[67,68]. In our study CAP have decreased the zeta potential of PVA-AgNP, which may be associated with absence of repulsive forces at the double layer leading to the likelihood of agglomeration that can be seen in uptake of PVA-AgNP in figure 5. Another reason for the agglomeration of NPs evident in the uptake of PVA-AgNP with CAP in figure 5a is due to serum proteins in culture media absorbed on NPs surface. Studies have reported the nanoparticle protein corona adds complexity to biological system interactions that cannot be limited to electrostatic binding alone. The new biological identity of the nanoparticle influences cell behaviour interaction[69–71].

Nanoparticle detection in cells and tissues are often achieved by employing electron microscopy techniques and confocal microscopy to investigate translocation of NPs in cells[72–74]. While these techniques have extensively accomplished identification of NPs, they lack the potential to validate NPs presence in cells or tissue via spectral mapping. The spectral imaging technique provides each pixel of SI image a spectral response for each spatial area of a pixel[75]. SI of NPs in cells provides the feasibility of detecting NPs, partial size, surface modification, spatial location, presence of NP agglomeration and wavelength differentiation[76,77]. In this study, we used SI to asses uptake of PVA-AgNP when exposed to CAP resulting to an enhanced PVA-AgNP uptake when cells are exposed to CAP than of NP treatment alone. This was quantified using atomic absorption spectroscopy where we confirmed a 50-fold increase in Ag/cell following CAP treatment. Our group demonstrated the direct and indirect chemical effects generated by CAP DIT-120 is a mediator of the uptake increase of AuNPs[29]. Our data here provide further evidence that CAP DIT-120 can stimulate uptake of nanomaterials of different sizes and compositions in addition to significantly enhancing cytotoxicity. Interestingly, Au/cell is increased by 50% following exposure to CAP, whereas Ag/cell is increased 50-fold under similar conditions. This may be due to nanomaterial size, the direct effect of CAP on the AgNP or due to a cellular process. Further investigation of the biological processes that regulate CAP-stimulated uptake of nanomaterials and cytotoxicity in GBM are ongoing and will offer future insights into adapting these combinational approaches for development of therapeutics for treatment of GBM and other solid tumours.

In conclusion, the current study reports the enhanced synergistic cytotoxic effect of the combined PVA-AgNP and CAP on U373MG cells *in vitro.* The study showed the ROS-dependent toxicity of the combined therapy, which was prevented by NAC. Enhanced uptake of PVA-AgNP followed by CAP treatment was confirmed using spectral imaging and AAS. Overall, the results indicate the effect of CAP on the physical properties of PVA-AgNP leading to decrease in nanoparticle size, decrease in surface charge distribution and inducing enhanced uptake, aggregation and synergistic cytotoxicity. The findings in the study demonstrated the combination therapy of PVA-AgNP with CAP can be further evaluated for its potential use in cancer therapy.

## Acknowledgements

This research work is supported by DIT Fiosraigh Research Scholarship programme (E.M.), European Research Council Starting Grant Number ERC-SG-335508 (A.G.), Science Foundation Ireland Grant Number 11/PI/08 (A.C.), Science Foundation Ireland Grant Number 14/IA/2626 (P.C. and J.C.), The authors also thank the FOCAS Research Institute, TU Dublin and UCD for the use of facilities.

## Author Contributions

E.M, A.C, P.C. and J.F.C. conceived the project and designed the experiments. E.M., A.G., and A.L. performed the experiments, collected and analysed the data. E.M., A.G., and J.F.C. co-wrote the paper. All authors discussed the results and reviewed the manuscript.

