## Supplementary Materials for "Cold Atmospheric Plasma induces silver nanoparticle uptake, oxidative dissolution and enhanced cytotoxicity in Glioblastoma multiforme cells"

**Supplemental Materials 1**

**Table of data for figure 1b**

| **Wavelength** | **Initial** | **3 months** | **6 months** |
| --- | --- | --- | --- |
| 350 | 0.2205 | 0.2311 | 0.2217 |
| 352 | 0.222 | 0.2329 | 0.2235 |
| 354 | 0.2239 | 0.2352 | 0.2257 |
| 356 | 0.226 | 0.2379 | 0.2282 |
| 358 | 0.2284 | 0.2408 | 0.2309 |
| 360 | 0.2311 | 0.2437 | 0.234 |
| 362 | 0.2342 | 0.2471 | 0.2375 |
| 364 | 0.2376 | 0.2512 | 0.2415 |
| 366 | 0.2427 | 0.2574 | 0.247 |
| 368 | 0.2486 | 0.2639 | 0.2538 |
| 370 | 0.2562 | 0.2723 | 0.2618 |
| 372 | 0.2642 | 0.2817 | 0.2707 |
| 374 | 0.2726 | 0.2918 | 0.2799 |
| 376 | 0.2808 | 0.3006 | 0.2887 |
| 378 | 0.2888 | 0.3097 | 0.2981 |
| 380 | 0.2966 | 0.3185 | 0.3063 |
| 382 | 0.3054 | 0.3287 | 0.316 |
| 384 | 0.3139 | 0.3382 | 0.3252 |
| 386 | 0.3235 | 0.3479 | 0.3358 |
| 388 | 0.3335 | 0.3586 | 0.3463 |
| 390 | 0.3426 | 0.3685 | 0.356 |
| 392 | 0.3503 | 0.3767 | 0.3643 |
| 394 | 0.3553 | 0.3824 | 0.3699 |
| 396 | 0.3591 | 0.3869 | 0.3742 |
| 398 | 0.3611 | 0.3894 | 0.3762 |
| 400 | 0.3609 | 0.3891 | 0.376 |
| 402 | 0.3586 | 0.3869 | 0.3736 |
| 404 | 0.3553 | 0.383 | 0.3702 |
| 406 | 0.3501 | 0.3773 | 0.3649 |
| 408 | 0.3436 | 0.3704 | 0.3579 |
| 410 | 0.3358 | 0.3616 | 0.3504 |
| 412 | 0.3264 | 0.3516 | 0.3408 |
| 414 | 0.3145 | 0.3388 | 0.3282 |
| 416 | 0.3022 | 0.3251 | 0.3151 |
| 418 | 0.2906 | 0.3123 | 0.3034 |
| 420 | 0.2775 | 0.2983 | 0.2891 |
| 422 | 0.2663 | 0.2863 | 0.2779 |
| 424 | 0.2557 | 0.2739 | 0.2666 |
| 426 | 0.2446 | 0.2625 | 0.2549 |
| 428 | 0.2347 | 0.2509 | 0.2444 |
| 430 | 0.2229 | 0.2391 | 0.2319 |
| 432 | 0.2107 | 0.2259 | 0.2196 |
| 434 | 0.2008 | 0.2149 | 0.2088 |
| 436 | 0.1907 | 0.2043 | 0.1982 |
| 438 | 0.1801 | 0.1929 | 0.1873 |
| 440 | 0.171 | 0.1825 | 0.1777 |
| 442 | 0.1627 | 0.1731 | 0.1689 |
| 444 | 0.1547 | 0.165 | 0.1607 |
| 446 | 0.1463 | 0.1559 | 0.1519 |
| 448 | 0.1395 | 0.1488 | 0.1452 |
| 450 | 0.1343 | 0.1426 | 0.1392 |

**Table of data for figure 1c**

| **Control** | | | **+6 months** | | |
| --- | --- | --- | --- | --- | --- |
| **Mean** | **SD** | **N** | **Mean** | **SD** | **N** |
| 10.68 | 1.98 | 3 | 9.59 | 2.52 | 3 |

**Supplemental Materials 2**

**Table of data for figure 2a**

|  | **Control Plate** | | | | | **5s CAP** | | | | | **10s CAP** | | | | | **25s CAP** | | | | | **40s CAP** | | | | |
| --- | --- | --- | --- | --- | --- | --- | --- | --- | --- | --- | --- | --- | --- | --- | --- | --- | --- | --- | --- | --- | --- | --- | --- | --- | --- |
| **0** | 91.68 | 95.45 | 94.05 | 98.58 | 98.09 | 91.11 | 97.37 | 90.66 | 93.81 | 91.27 | 94.89 | 91.30 | 91.12 | 92.37 | 94.77 | 89.26 | 87.71 | 82.22 | 85.20 | 84.71 | 72.05 | 77.88 | 82.22 | 87.66 | 87.17 |
| **0.008** | 86.16 | 87.63 | 93.84 | 95.02 | 92.33 | 87.50 | 95.32 | 83.82 | 79.79 | 84.56 | 84.25 | 84.37 | 82.50 | 87.44 | 82.61 | 63.77 | 57.81 | 65.28 | 68.20 | 61.22 | 49.61 | 54.61 | 53.16 | 47.05 | 60.10 |
| **0.024** | 89.76 | 81.04 | 85.63 | 89.21 | 90.88 | 86.44 | 87.21 | 79.44 | 78.00 | 83.35 | 78.92 | 84.01 | 89.00 | 84.94 | 87.98 | 55.96 | 58.10 | 51.70 | 58.38 | 61.38 | 43.73 | 46.97 | 48.92 | 42.30 | 40.02 |
| **0.07** | 88.11 | 89.00 | 83.55 | 86.81 | 91.66 | 83.66 | 83.19 | 75.43 | 77.74 | 79.92 | 81.20 | 81.53 | 80.60 | 86.04 | 72.68 | 48.64 | 52.73 | 49.81 | 49.45 | 52.55 | 26.72 | 25.53 | 23.22 | 30.86 | 22.96 |
| **0.22** | 86.86 | 88.89 | 83.51 | 89.58 | 80.35 | 86.19 | 82.39 | 83.37 | 73.76 | 75.67 | 81.29 | 83.93 | 81.05 | 76.91 | 74.69 | 47.88 | 47.25 | 49.53 | 47.70 | 45.82 | 23.04 | 25.43 | 21.69 | 24.09 | 24.16 |
| **0.64** | 89.60 | 85.28 | 85.50 | 83.63 | 87.00 | 79.34 | 83.72 | 82.52 | 84.11 | 76.51 | 78.51 | 79.73 | 80.06 | 79.34 | 75.93 | 40.83 | 39.15 | 41.91 | 44.19 | 41.59 | 21.70 | 18.53 | 24.52 | 23.83 | 22.19 |
| **1.95** | 77.65 | 83.74 | 76.15 | 75.04 | 75.16 | 78.39 | 74.02 | 73.73 | 76.09 | 76.37 | 78.43 | 71.55 | 73.73 | 79.89 | 77.60 | 26.25 | 35.26 | 25.51 | 27.88 | 33.41 | 17.50 | 12.89 | 19.53 | 15.38 | 14.21 |
| **5.84** | 54.30 | 51.41 | 53.19 | 55.07 | 46.95 | 51.80 | 59.05 | 52.66 | 59.41 | 62.00 | 56.19 | 61.93 | 58.77 | 53.71 | 60.83 | 13.58 | 16.63 | 12.36 | 19.50 | 16.33 | 8.77 | 8.45 | 10.63 | 13.19 | -0.14 |
| **17.54** | -0.15 | -0.45 | -0.21 | -0.07 | -0.19 | -0.32 | -0.38 | -0.30 | -0.17 | -0.27 | -0.29 | -0.30 | -0.01 | -0.18 | -0.20 | -0.18 | -0.16 | -0.25 | -0.20 | -0.19 | -0.32 | 0.05 | -0.11 | -0.24 | -0.20 |

**Table of data for figure 2b**

| **Dose AgNP** | **Dose CAP** | **Effect** | **Combination Index** |
| --- | --- | --- | --- |
| 4.09 | 5 | 0.5 | 0.93 |
| 3.95 | 10 | 0.5 | 0.96 |
| 0.079 | 25 | 0.5 | 0.35 |
| 0.01 | 40 | 0.5 | 0.54 |

**Supplemental Materials 3**

**Table of data for figure 3a**

|  | **AgNP Control** | | | | | **AgNP + 25s CAP** | | | | | **AgNP + NAC** | | | | | **AgNP + NAC + 25s CAP** | | | | |
| --- | --- | --- | --- | --- | --- | --- | --- | --- | --- | --- | --- | --- | --- | --- | --- | --- | --- | --- | --- | --- |
| **0** | 101.26 | 100.54 | 97.57 | 99.88 | 99.35 | 75.13 | 74.61 | 78.83 | 80.80 | 76.45 | 84.58 | 101.32 | 100.31 | 98.53 | 84.63 | 87.38 | 87.61 | 86.64 | 76.91 | 75.76 |
| **0.008** | 98.23 | 95.70 | 96.61 | 93.73 | 95.66 | 61.35 | 73.86 | 59.44 | 56.09 | 55.55 | 99.09 | 93.58 | 97.12 | 93.62 | 92.62 | 69.42 | 68.84 | 62.89 | 67.74 | 60.63 |
| **0.024** | 92.12 | 93.03 | 94.13 | 95.81 | 88.27 | 47.73 | 49.02 | 53.15 | 51.40 | 50.22 | 82.19 | 92.00 | 81.20 | 90.57 | 94.44 | 66.93 | 70.68 | 80.41 | 69.03 | 65.79 |
| **0.072** | 83.15 | 82.25 | 97.86 | 81.32 | 85.57 | 49.96 | 59.87 | 48.72 | 55.76 | 45.05 | 83.62 | 84.26 | 77.41 | 80.88 | 88.27 | 76.99 | 79.49 | 80.35 | 68.40 | 78.20 |
| **0.22** | 83.04 | 78.97 | 84.22 | 92.71 | 80.85 | 42.24 | 42.45 | 51.16 | 51.41 | 48.37 | 82.05 | 76.34 | 94.34 | 100.65 | 92.00 | 62.92 | 61.05 | 79.34 | 63.98 | 75.49 |
| **0.64** | 89.21 | 75.00 | 81.86 | 71.06 | 82.78 | 36.99 | 37.28 | 39.55 | 49.754 | 49.15 | 86.88 | 83.34 | 88.10 | 81.68 | 88.85 | 61.35 | 60.16 | 68.86 | 73.55 | 70.93 |
| **1.95** | 82.15 | 85.63 | 81.11 | 80.48 | 88.50 | 35.89 | 32.58 | 48.48 | 33.07 | 40.66 | 86.72 | 90.85 | 88.73 | 93.45 | 81.94 | 72.62 | 69.91 | 63.96 | 56.37 | 69.23 |
| **5.84** | 49.73 | 45.49 | 63.52 | 39.49 | 51.91 | 9.94 | 6.54 | 6.60 | 14.28 | 10.85 | 62.82 | 61.59 | 66.97 | 55.30 | 60.02 | 36.35 | 35.14 | -0.28 | -0.12 | -0.03 |
| **17.54** | -0.24 | -0.41 | -0.32 | -0.19 | -0.33 | -0.28 | -0.06 | -0.22 | -0.41 | -0.33 | -0.07 | -0.20 | -0.13 | -0.26 | -0.22 | -0.29 | -0.28 | -0.23 | -0.20 | -0.08 |

**Supplemental Materials 4**

**Table of data for figure 4a**

| **CAP (sec)** | **PVA-AgNP in reaction solution (including NaBH4)** | | | **Purified PVA-AgNP** | | | **Purified PVA-AgNP resuspended in 4mM NaBH4** | | |
| --- | --- | --- | --- | --- | --- | --- | --- | --- | --- |
|  | **Mean** | **SD** | **N** | **Mean** | **SD** | **N** | **Mean** | **SD** | **N** |
| **0** | 8.45 | 1.856 | 9 | 8.83 | 1.02 | 6 | 8.764 | 2.197 | 3 |
| **5** | 6.77 | 1.11 | 6 |  |  |  |  |  |  |
| **10** | 6.22 | 1.07 | 9 | 8.804 | 1.01 | 6 | 7.894 | 2.08 | 3 |
| **25** | 3.94 | 1.09 | 6 | 8.823 | 1.22 | 6 | 4.646 | 1.043 | 3 |
| **40** | 2.29 | 1.02 | 9 | 8.925 | 1.046 | 6 | 4.074 | 1.042 | 3 |
| **80** | 1.52 | 0.62 | 9 | 8.78 | 1.1 | 6 | 3.265 | 0.784 | 3 |

**Table of data for figure 4c**

| **PVA-AgNP in mH_2_O** | | | **PVA-AgNP in mH_2_O + CAP** | | |
| --- | --- | --- | --- | --- | --- |
| **Mean** | **SD** | **N** | **Mean** | **SD** | **N** |
| -55.6 | 8.06 | 3 | -26.8 | 9.08 | 3 |

| **PVA-AgNP in DMEM** | | | **PVA-AgNP in DMEM + CAP** | | |
| --- | --- | --- | --- | --- | --- |
| **Mean** | **SD** | **N** | **Mean** | **SD** | **N** |
| -18.00 | 8.70 | 3 | -11.40 | 5.06 | 3 |

**Supplemental Materials 5**

**Table of data for figure 5a**

```
%%supplemental functions available from second author on request

%read spectral image 'cells.agnp.cap39_4.9'
Img_NP=eraCV('cells.agnp.cap39_4.9');
WL_VIS=[399.7643, 400.9560, 402.1481, 403.3406, 404.5335,405.7268, 406.9205, 408.1147, 409.3092, 410.5042,411.6996, 412.8954, 414.0916, 415.2882, 416.4853,417.6827, 418.8806, 420.0789, 421.2776, 422.4767,423.6762, 424.8762, 426.0765, 427.2773, 428.4785,429.6801, 430.8821, 432.0845, 433.2874, 434.4906,435.6943, 436.8984, 438.1029, 439.3078, 440.5131,441.7189, 442.9250, 444.1316, 445.3386, 446.5460,447.7538, 448.9620, 450.1707, 451.3797, 452.5892,453.7991, 455.0094, 456.2201, 457.4312, 458.6428,459.8547, 461.0671, 462.2799, 463.4931, 464.7067,465.9207, 467.1351, 468.3500, 469.5653, 470.7809,471.9971, 473.2136, 474.4305, 475.6478, 476.8656,478.0838, 479.3024, 480.5214, 481.7408, 482.9606,484.1808, 485.4015, 486.6226, 487.8441, 489.0659,490.2882, 491.5110, 492.7341, 493.9577, 495.1816,496.4060, 497.6308, 498.8560, 500.0817, 501.3077,502.5341, 503.7610, 504.9883, 506.2160, 507.4441,508.6726, 509.9016, 511.1309, 512.3607, 513.5909,514.8215, 516.0525, 517.2839, 518.5157, 519.7480,520.9807, 522.2137, 523.4473, 524.6812, 525.9155,527.1503, 528.3854, 529.6210, 530.8569, 532.0933,533.3301, 534.5674, 535.8050, 537.0430, 538.2815,539.5204, 540.7597, 541.9994, 543.2395, 544.4801,545.7210, 546.9624, 548.2042, 549.4464, 550.6890,551.9320, 553.1754, 554.4193, 555.6636, 556.9083,558.1533, 559.3988, 560.6448, 561.8911, 563.1378,564.3850, 565.6326, 566.8806, 568.1290, 569.3778,570.6271, 571.8767, 573.1268, 574.3773, 575.6282,576.8795, 578.1312, 579.3833, 580.6359, 581.8889,583.1422, 584.3960, 585.6502, 586.9048, 588.1599,589.4153, 590.6712, 591.9275, 593.1842, 594.4413,595.6989, 596.9568, 598.2151, 599.4739, 600.7330,601.9927, 603.2527, 604.5131, 605.7739, 607.0352,608.2968, 609.5589, 610.8214, 612.0843, 613.3477,614.6113, 615.8755, 617.1400, 618.4050, 619.6704,620.9363, 622.2025, 623.4691, 624.7362, 626.0037,627.2715, 628.5398, 629.8085, 631.0776, 632.3472,633.6171, 634.8875, 636.1583, 637.4294, 638.7010,639.9731, 641.2455, 642.5183, 643.7916, 645.0653,646.3394, 647.6139, 648.8888, 650.1642, 651.4399,652.7161, 653.9927, 655.2697, 656.5470, 657.8248,659.1030, 660.3817, 661.6608, 662.9403, 664.2202,665.5005, 666.7812, 668.0624, 669.3439, 670.6259,671.9082, 673.1910, 674.4742, 675.7578, 677.0419,678.3263, 679.6112, 680.8965, 682.1821, 683.4683,684.7548, 686.0417, 687.3290, 688.6168, 689.9050,691.1936, 692.4825, 693.7720, 695.0618, 696.3521,697.6427, 698.9338, 700.2253, 701.5172, 702.8095,704.1023, 705.3954, 706.6890, 707.9830, 709.2773,710.5721, 711.8674, 713.1630, 714.4590, 715.7555,717.0524, 718.3496, 719.6473, 720.9455, 722.2440,723.5430, 724.8423, 726.1421, 727.4423, 728.7429,730.0439, 731.3453, 732.6472, 733.9495, 735.2521,736.5552, 737.8587, 739.1626, 740.4669, 741.7717,743.0769, 744.3824, 745.6884, 746.9948, 748.3016,749.6089, 750.9165, 752.2246, 753.5331, 754.8419,756.1512, 757.4609, 758.7711, 760.0817, 761.3926,762.7040, 764.0157, 765.3280, 766.6406, 767.9536,769.2671, 770.5809, 771.8952, 773.2098, 774.5250,775.8405, 777.1564, 778.4728, 779.7896, 781.1067,782.4243, 783.7423, 785.0607, 786.3795, 787.6987,789.0184, 790.3385, 791.6590, 792.9799, 794.3012,795.6229, 796.9451, 798.2676, 799.5906, 800.9140,802.2378, 803.5620, 804.8866, 806.2117, 807.5372,808.8630, 810.1893, 811.5160, 812.8431, 814.1707,815.4986, 816.8270, 818.1558, 819.4849, 820.8145,822.1445, 823.4750, 824.8058, 826.1371, 827.4688,828.8008, 830.1333, 831.4662, 832.7996, 834.1333,835.4674, 836.8020, 838.1370, 839.4724, 840.8082,842.1445, 843.4811, 844.8181, 846.1556, 847.4935,848.8318, 850.1705, 851.5096, 852.8492, 854.1891,855.5295, 856.8703, 858.2115, 859.5531, 860.8951,862.2375, 863.5804, 864.9237, 866.2673, 867.6115,868.9560, 870.3009, 871.6462, 872.9920, 874.3382,875.6848, 877.0317, 878.3792, 879.7271, 881.0753,882.4239, 883.7729, 885.1224, 886.4724, 887.8226,889.1733, 890.5245, 891.8761, 893.2280, 894.5804,895.9332, 897.2865, 898.6401, 899.9941, 901.3486,902.7035, 904.0587, 905.4144, 906.7705, 908.1271,909.4840, 910.8413, 912.1991, 913.5573, 914.9159,916.2749, 917.6343, 918.9941, 920.3544, 921.7151,923.0762, 924.4376, 925.7996, 927.1619, 928.5247,929.8878, 931.2514, 932.6154, 933.9797, 935.3445,936.7098, 938.0754, 939.4415, 940.8080, 942.1748,943.5421, 944.9099, 946.2780, 947.6465, 949.0154,950.3848, 951.7546, 953.1248, 954.4954, 955.8665,957.2379, 958.6097, 959.9820, 961.3547, 962.7278,964.1013, 965.4752, 966.8495, 968.2243, 969.5995,970.9750, 972.3510, 973.7274, 975.1042, 976.4815,977.8591, 979.2372, 980.6157, 981.9946, 983.3739,984.7537, 986.1338, 987.5143, 988.8953, 990.2767,991.6584, 993.0406, 994.4233, 995.8063, 997.1898,998.5736, 999.6118];
%read RGB image for 'cells.agnp.cap39_4.9'
Img_NP_RGB=imread('cells.agnp.cap39_4.9_RGB.png');

%select coordinates from
[Coord]=Region_select(Img_NP_RGB);

%extract selected spectra from each class in SpecIm
SpecIm.Y=[];
SpecIm.X=[];

for i=1:size(Coord,2)
for j=1:size(Coord{1,i}.coords,1)
SpecIm.Y=[SpecIm.Y;sprintf(('''%s'','),Coord{1,i}.label)];
SpecIm.X=[SpecIm.X;squeeze(Img_NP(Coord{1,i}.coords(j,2),Coord{1,i}.coords(j,1),:))'];
end
end

%Make Figure 5a
cols=['g';'b';'r';'c';'m';'y'];
figure,imshow(Img_NP_RGB)
for i=2:size(Coord,2)
    for j=1:size(Coord{1,i}.coords,1)
hold on,plot(Coord{1,i}.coords(j,1),Coord{1,i}.coords(j,2),'+', 'MarkerSize', 10,'LineWidth',2,'Color',cols(i))
    end
end

%note Coordinates selected for Figure 5a were:
```

| **Cell coordinates** | | **NP coordinates** | |
| --- | --- | --- | --- |
| **Row** | **Column** | **Row** | **Column** |
| 404 | 231 | 299 | 190 |
| 408 | 240 | 280 | 149 |
| 412 | 260 | 292 | 163 |
| 412 | 281 | 319 | 151 |
| 403 | 307 | 297 | 146 |
| 394 | 334 | 296 | 133 |
| 386 | 365 | 331 | 166 |
| 356 | 356 | 357 | 160 |
| 329 | 355 | 360 | 194 |
| 257 | 446 | 330 | 203 |
| 291 | 463 | 201 | 567 |
| 315 | 488 | 216 | 583 |
| 333 | 557 | 230 | 598 |
| 327 | 582 | 238 | 614 |
| 284 | 631 | 239 | 626 |
| 269 | 647 | 244 | 645 |
| 201 | 640 |  |  |
| 175 | 604 |  |  |
| 154 | 588 |  |  |
| 145 | 520 |  |  |
| 197 | 446 |  |  |
| 197 | 397 |  |  |
| 258 | 136 |  |  |
| 257 | 166 |  |  |

%Pool the spectra and apply standard normal variate pretreatment

X_NCC=[jsnv((SpecIm.X(SpecIm.Y(:,2)=='N',:)));jsnv((SpecIm.X(SpecIm.Y(:,2)=='C',:)))];

Y_NCC=[1*ones(size(SpecIm.Y(SpecIm.Y(:,2)=='N'),1),1);2*ones(size(SpecIm.Y(SpecIm.Y(:,2)=='C'),1),1)];

%plot the spectra

figure,plot(WL_VIS,mean(X_NCC(Y_NCC==1,:)),'r')

hold on,plot(WL_VIS,mean(X_NCC(Y_NCC==2,:)),'b')

xlabel('Wavelength (nm)')

ylabel('SNV(Reflectance)')

**Supplementary graph for figure 5b**

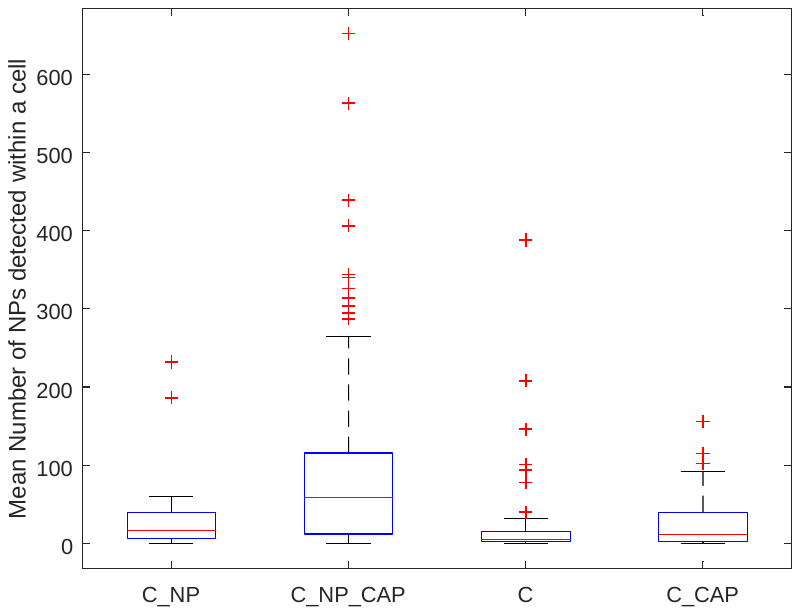

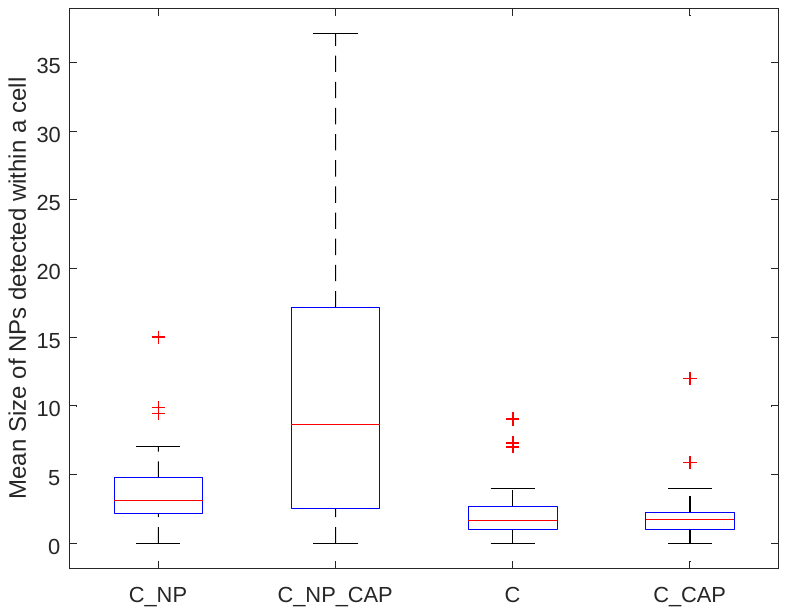

```
%count the nps
for I=1:11
    for J=1:num_im(I)
        cc_re=Pred_NP{I,J}.cc;
        cc_re=bwareaopen(cc_re,300);
        cc = bwlabel(cc_re);
        outstats = regionprops(cc_re,'Area','Circularity','Eccentricity');
        for i=1:size(outstats,1)
            %get shape and size of cell
            Mask_new=cc==i;

            %get nanos in Mask new
            Mask_NP=Mask_new.*Pred_NP{I,J}.Img_RGB_af(:,:,1);
            outstatsNP = bwconncomp(Mask_NP);
            APred_NP{I,J}.CellP{i}=[];
            APred_NP{I,J}.CellP{i}.NumNP=size(outstatsNP.PixelIdxList,2);
            APred_NP{I,J}.CellP{i}.Size_Shape=outstats(i);

            if size(outstatsNP.PixelIdxList,2)>0
                for j=1:size(outstatsNP.PixelIdxList,2)
                    APred_NP{I,J}.CellP{i}.Size_NP(j)=numel(outstatsNP.PixelIdxList{1,j});
                end
                APred_NP{I,J}.CellP_Im{i}=0.3*Mask_new+Mask_NP;
            else
                APred_NP{I,J}.CellP{i}.Size_NP=0;
                APred_NP{I,J}.CellP{i}.NumNP=0;
                APred_NP{I,J}.CellP{i}.Size_Shape=0;
            end

        end
    end
end

%extract # NPs per cell
Num_NP=[];
Size_NP=[];
Size_Cell=[];
Circularity_Cell=[];
Eccentricity_Cell=[];

Y=[];
for I=1:11

        for J=1:num_im(I)

            for i = 1:size(APred_NP{I,J}.CellP,2)

            Num_NP=[Num_NP;APred_NP{I,J}.CellP{i}.NumNP];
            Size_NP=[Size_NP;mean(APred_NP{I,J}.CellP{i}.Size_NP)];
            if APred_NP{I,J}.CellP{i}.Size_NP>0
            Size_Cell=[Size_Cell;APred_NP{I,J}.CellP{i}.Size_Shape.Area];
            Circularity_Cell=[Circularity_Cell;APred_NP{I,J}.CellP{i}.Size_Shape.Circularity];
            Eccentricity_Cell=[Eccentricity_Cell;APred_NP{I,J}.CellP{i}.Size_Shape.Eccentricity];
            else
               Size_Cell=[Size_Cell;0];
            Circularity_Cell=[Circularity_Cell;0];
            Eccentricity_Cell=[Eccentricity_Cell;0];
            end

            Y=[Y;[I,J,i]];

            end
        end
end

treatment={'C_NP';'C_NP_CAP';'C_NP';'C_NP_CAP';'C';'C_CAP';'C_CAP';'C_NP_CAP';'C_NP_CAP';'C_NP_CAP';'C'};

%make a table of all results
for i=1:size(Y,1)
Y_tr(i)=treatment(Y(i,1));
end

T=table(Y(:,1),Y(:,2),Y(:,3),Y_tr',Num_NP,Size_NP,Size_Cell,Circularity_Cell,'VariableNames',{'I','J','Cell','Treatment','Num_NP','Size_NP','Size_Cell','Circularity_Cell'});

%remove the doubles

T_remove_doubles=T([1:86,88:90,92:103,105:234,236:end],:);

n_C_NP=find(strcmp(T_remove_doubles.Treatment, 'C_NP'));
n_C_NP_CAP=find(strcmp(T_remove_doubles.Treatment, 'C_NP_CAP'));
n_C=find(strcmp(T_remove_doubles.Treatment, 'C'));
n_C_CAP=find(strcmp(T_remove_doubles.Treatment, 'C_CAP'));

%median of each measurement
%Box plot
figure,set(gca,'Color',[1,1,1]),boxplot(T_remove_doubles.Num_NP,T_remove_doubles.Treatment)
ylabel('Mean Number of NPs detected within a cell')

figure,boxplot(T_remove_doubles.Size_NP,T_remove_doubles.Treatment)
ylabel('Mean Size of NPs detected within a cell')

figure,boxplot(T_remove_doubles.Size_Cell,T_remove_doubles.Treatment)
ylabel('Size of cell')

figure,boxplot(T_remove_doubles.Circularity_Cell,T_remove_doubles.Treatment)
ylabel('Circularity of cell')
```

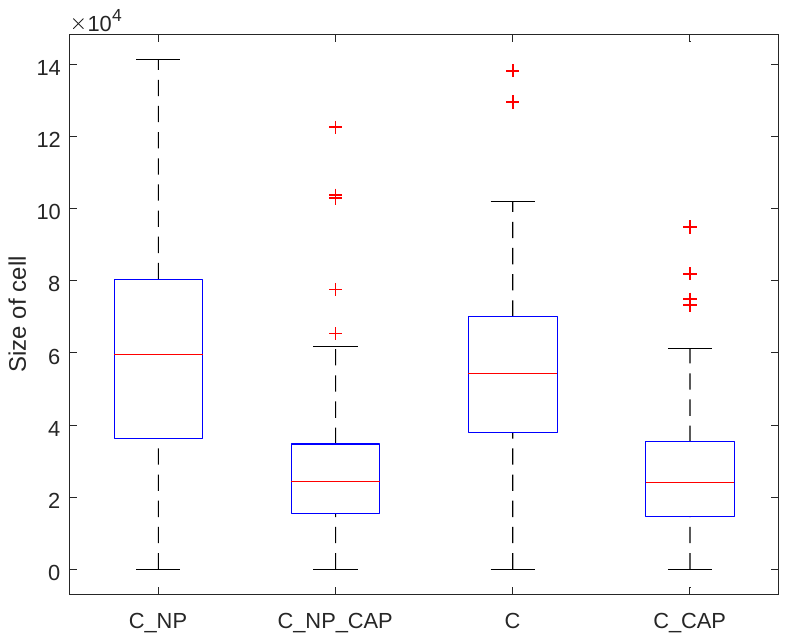

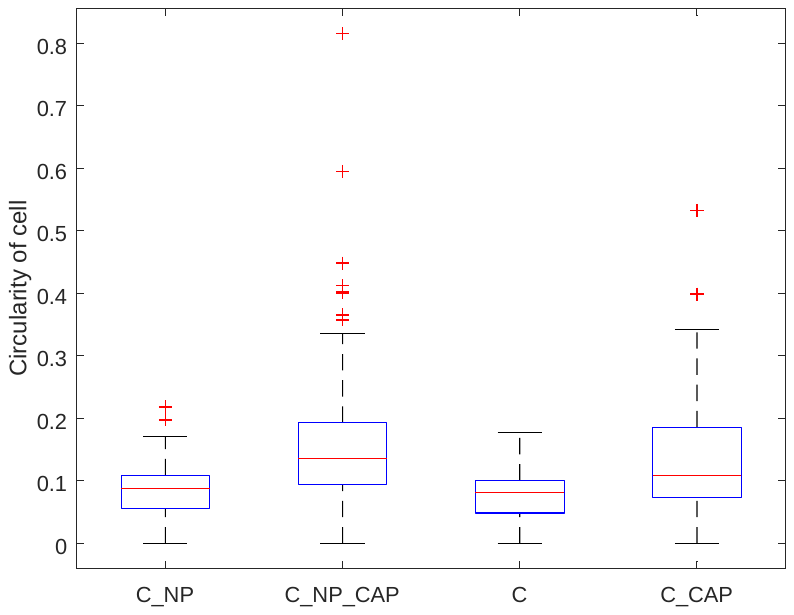

**Table of data for figure 5c**

| **Negative Control** | | | | **PVA-AgNP** | | | | **CAP** | | | | **PVA-AgNP + 25s CAP** | | | |
| --- | --- | --- | --- | --- | --- | --- | --- | --- | --- | --- | --- | --- | --- | --- | --- |
| **0.013** | **0** | **-0.0198** | **-0.007** | **0.01904** | **0.0118** | **0.0116** | **0.08** | **0** | **0** | **0** | **0** | **0.8563** | **1.0852** | **1.06** | **0.55** |
